# Model-X knockoffs reveal data-dependent limits on regulatory network identification

**DOI:** 10.1101/2023.05.23.541948

**Authors:** Eric Kernfeld, Rebecca Keener, Patrick Cahan, Alexis Battle

## Abstract

Computational biologists have long sought to automatically infer transcriptional regulatory networks (TRNs) from gene expression data, but such approaches notoriously suffer from false positives. Two points of failure could yield false positives: faulty hypothesis testing, or erroneous assumption of a classic criterion called *causal sufficiency*. We show that a recent statistical development, model-X knockoffs, can effectively control false positives in tests of conditional independence in mouse and *E. coli* data, which rules out faulty hypothesis tests. Yet, benchmarking against ChIP and other gold standards reveals highly inflated false discovery rates. This identifies the causal sufficiency assumption as a key limiting factor in TRN inference.

## Background

The prospect of reliably determining transcriptional regulatory networks (TRNs) from gene expression measurements has been enticing since the advent of gene expression profiling (Duggan, Bittner, Chen, Meltzer, & Trent, 1999; Liang, Fuhrman, & Somogyi, 1998). We define a TRN as a graph where edges connect transcription factors (TF’s) to target genes that they directly regulate. To the extent that such networks are accurate, they would enable systems-based approaches to study complex biological processes. Examples of applications that accurate TRNs would enable include: predicting the effects of genetic perturbations during differentiation and development (Kamimoto et al., 2023), revealing genetic architecture of complex traits (Boyle, Li, & Pritchard, 2017; Freimer et al., 2022; Krishnan et al., 2016); and reverse engineering transcriptional changes to aid in drug development for cancer (Baca et al., 2021) (Reddy et al., 2021), aging (Lee et al., 2021), and heart disease (Amrute et al., 2022). These are just a few of the many applications across diverse fields of biological research in which accurate TRN models would yield useful advances.

Given the huge potential of accurate TRNs, dozens of TRN inference methods have been invented (reviewed in (Nguyen, Tran, Tran, Pehlivan, & Nguyen, 2021) and (Sanguinetti & Huynh-Thu, 2019)). It is reassuring that some TRNs inferred using these methods are enriched for functionally relevant gene pairs or sets (e.g. (Parsana et al., 2019), (Margolin et al., 2006), (Chasman et al., 2019), (Cote, Young, & Huckins, 2022; Morgan, Tjärnberg, Nordling, & Sonnhammer, 2019), (Diaz & Stumpf, 2022)), but this enrichment does not guarantee the reliability of individual regulatory hypotheses. In fact, the best performing TRN inference methods showed early precision around ∼50% in a seminal *E. coli* benchmarking project (Marbach et al., 2012), and in recent benchmarks on mammalian data, early precision with respect to cell-type specific ChIP data is at most 1.7 times better than random (Pratapa, Jalihal, Law, Bharadwaj, & Murali, 2020). Therefore, a key obstacle to realizing the potential of TRN inference methods is the risk of false positives (Diaz & Stumpf, 2022).

False discovery rate (FDR) control is a way to cull conclusions from experiments that yield thousands of statistical hypothesis tests by estimating the expected proportion of false positives among the significant findings (Benjamini & Hochberg, 1995). FDR control has become standard in many fields, including differential gene expression analysis (Korthauer et al., 2019) and neuroimaging (Genovese, Lazar, & Nichols, 2002), due to its simple interpretation and useful balance between stringency and power (Benjamini & Hochberg, 1995). While TRN methods would similarly benefit from FDR control, there are unique challenges to achieving FDR control in this context, as exemplified by the TRN inference methods that attempt it. These fall into two broad categories. One type forms a background distribution by randomly permuting expression levels of a given gene across samples (Chasman et al., 2019), (Kimura, Fukutomi, Tokuhisa, & Okada, 2020; Morgan et al., 2019). This strategy implies the following strong null hypothesis: *each gene is independent of all other genes unless it directly regulates them*. Thus, permutation tests will mistake indirect effects for direct effects (Barber & Candès, 2015); (Fithian & Lei, 2020) (Fig. 1A). Other FDR-controlled TRN inference methods do account for indirect effects, but they assume target transcript levels are a linear function of their regulators (Kim, 2015; Schäfer & Strimmer, 2005a) or have at most one regulator (Mukhopadhyay & Chatterjee, 2007). Kinetic models of transcription are not linear, and also not fully understood (Eck et al., 2020), so any method assuming linearity may incur excess false discoveries (Fig. 1A). An ideal method would require minimal or no assumptions about the quantitative relationship between each gene and its regulators. Finally, we note that popular TRN inference methods based on tree ensembles or mutual information can account for both nonlinearity and indirect relationships, but they do not provide finite-sample FDR control (Fig. 1A). Importantly, empirical tests of advertised FDR rates have not been reported for any category of TRN inference method.

**Figure 1.**
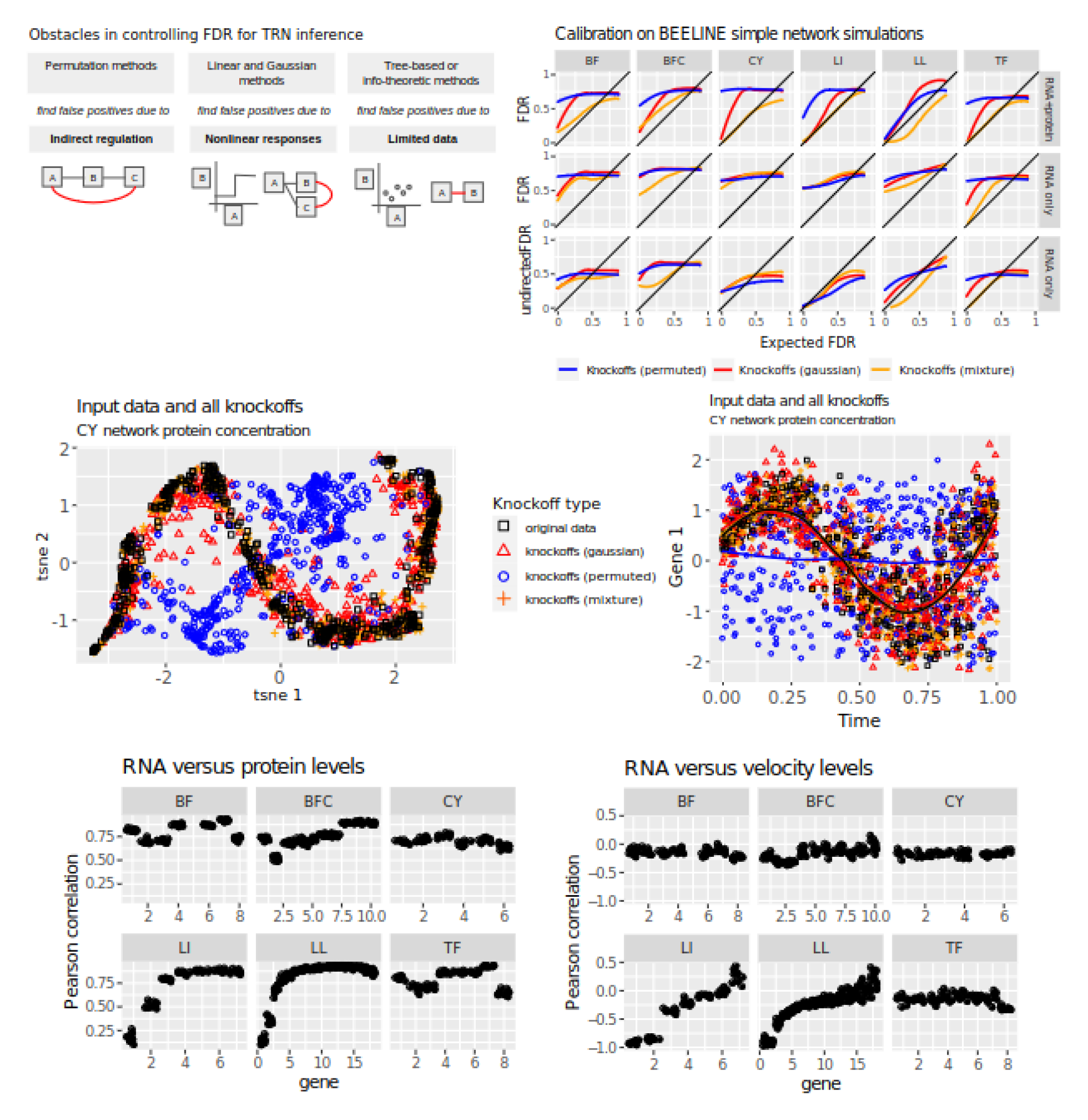
**Under ideal conditions, model-X knockoffs control FDR on a standard benchmark of network inference performance.** This figure contains only simulated data. **A)** Schematic depicting expected causes of false discoveries for permutation-based, Gaussian, and tree-based or information-theoretic TRN methods. **B)** Expected versus observed FDR for knockoff-based hypothesis tests used to nominate regulators in the toy networks from the BEELINE benchmarking framework (Pratapa et al. 2019). The six networks are listed across the top: bifurcating (BF), bifurcating converging (BFC), cyclic (CY), linear (LI), linear long (LL), and trifurcating (TF). In the bottom rows (“rna only”), RNA concentrations are used as input to the algorithm, following Pratapa et al. In the top row (“RNA + protein”), also revealed are protein concentrations and RNA production rates. Three methods of knockoff construction are used: independent permutation of all features (“permuted”), second-order knockoffs (“Gaussian”), and Gaussian mixture model knockoffs (“mixture”); (Gimenez et al. 2019). Results are averaged over 10 independent simulations. **C)** Protein expression for all genes in a realization of the “BFC” network (n=500), along with three types of knockoffs (n=500 each), all jointly reduced to two dimensions via t-Stochastic Neighbor Embedding (t-SNE)(van der Maaten & Hinton, 2008). **D)** Protein concentration and corresponding knockoff features for gene 1 in a realization of the “CY” network, plotted against time. No cell is measured twice; each dot is the terminus of an independent trajectory. Time is not used as input for generating knockoffs. Pearson correlation between RNA and protein concentration for each gene across all simulations used. **E)** Correlation between each TF and its knockoff when using Gaussian model-X knockoffs with sample covariance. **F)** Pearson correlation between RNA concentration and RNA production rate for each gene across all simulations used.

Part of the difficulty in empirical checks on FDR is that excess FDR could have a variety of causes. Of the assumptions required for causal inference (Spirtes, Glymour, & Scheines, 1993), two are most relevant to FDR control. First, conditional independence structure must be correctly inferred, which cannot be guaranteed when also assuming independence or linearity as discussed above. Second, all factors affecting transcript levels must be observed; this assumption is called *causal sufficiency*. It is unclear whether measuring mRNA levels of candidate regulators is enough to approximate causal sufficiency in a typical application of a TRN inference method, especially given the possibility of batch effects (Parsana et al., 2019). In principle, either type of obstacle could explain false discoveries generated by TRN inference methods, which limits the insight available even from careful real-data benchmarks.

In this work, we address these issues via recent statistical advances known as the *knockoff filter* and *model-X knockoffs* (supplementary file S1). The knockoff filter accounts for indirect effects without assuming all effects are linear. Importantly, knockoff-based FDR control on real data can be checked in a way that claims only conditional independence, without causal interpretation. We use such checks to determine whether the knockoff filter controls FDR in tests of conditional independence on mouse and *E. coli* datasets. We release open-source software to facilitate re-use of our methods for fast conditional independence testing (supplementary file S2).

Next, to improve causal interpretation of conditional independence findings, we devise a method to compare expected FDR from the knockoff filter against observed FDR on gold standards. This method yields unbiased results even when the gold standards are incomplete. Finally, we apply the entire pipeline to real data, producing high-confidence sets of conditional dependence relationships and comparing them to incomplete gold standards compiled from ChIP and knockout data. This provides the first empirical tests of FDR control and causal sufficiency on a real TRN task.

## Results

### In biochemical simulations, model-X knockoffs control FDR in TRN inference without using the true data-generating distribution

To test the reliability of model-X knockoffs in a controlled setting, we used the previously published simulated network data from the BEELINE TRN inference benchmarking framework (Pratapa et al., 2020). We generated knockoffs under three different modeling assumptions. The method labeled “Gaussian” used Gaussian knockoffs with the sample covariance matrix, which are simple to construct and may be adequate given the robustness properties of the knockoff filter (Barber, Candès, & Samworth, 2020). The method labeled “mixture” used a Gaussian mixture model (Gimenez, Ghorbani, & Zou, 2019), which provides more flexibility than a Gaussian for cases where the data are nonlinear or multimodal. The method labeled “permuted” randomly permuted samples within each gene (independent of the permutation applied to the other genes). Permutation implies very strict assumptions on the distribution of the features and is not expected to yield adequate knockoffs; we included it as an approximation of several existing permutation methods for error control in TRN inference (Chasman et al., 2019) (Kimura et al., 2020) (Morgan et al., 2019) (Verny, Sella, Affeldt, Singh, & Isambert, 2017). We provided the simulated data to the knockoff filter using only RNA expression levels (“RNA only”) or revealing RNA expression, RNA production rate, and protein levels (“RNA + protein”). With RNA only, static methods cannot infer directionality, so for RNA only, FDR was calculated with backwards edges counted as correct (Fig. 1B). Using networks capable of generating a variety of temporal trajectories, these experiments provide a baseline expectation for the behavior of the knockoff filter in TRN inference.

Results demonstrated that FDR control relies on two requirements: the assumed distribution of the regulators must match the true distribution closely, and causes of transcription must be completely observed (causal sufficiency). Regarding distributional assumptions, more flexible knockoffs improved FDR control, with the mixture model performing best, and with the cyclic network structure being especially difficult for Gaussian knockoffs to accommodate (Fig. 1B). Examples of knockoffs plotted against original data demonstrated obvious deficits of permuted knockoffs as negative controls, but only subtle weaknesses for Gaussian knockoffs, while mixture-model knockoffs were visually indistinguishable from the original data (Fig. 1C,D). Regarding causal sufficiency, this criterion was satisfied given RNA + protein data, which led to better FDR control, while RNA only did not satisfy causal sufficiency and had worse FDR control. Correlations between protein and RNA levels or RNA production rate and RNA levels were sometimes low or negative (Fig. 1E,F), and can also be poor in real data (de Sousa Abreu, Penalva, Marcotte, & Vogel, 2009). In principle, distributional assumptions and causal sufficiency are both important for error control in TRN inference, although in real datasets, either requirement may be limiting.

### Model-X knockoffs control FDR in testing conditional independence on a large, diverse *E. coli* dataset

To isolate the issue of distributional assumptions, we will use several diagnostics that test model-X knockoff performance on real data without requiring causal interpretation or external gold standards. We chose the Many Microbe Microarrays Database, which comprises gene expression data for 4,511 genes, including 334 transcription factors (TF), across 805 *E. coli* samples (Faith et al., 2008). As with the BEELINE data, knockoffs were constructed using a Gaussian distribution based on the sample covariance matrix; using a Gaussian mixture model (cluster assignments are shown in Fig. S1A); and using independent permutation of each gene. However, the Many Microbe Microarrays dataset is higher-dimensional than the BEELINE data and the sample covariance matrix may be a poor estimator (Schäfer & Strimmer, 2005b). Therefore, we tested four additional sets of Gaussian knockoffs based on established methods for high-dimensional covariance estimation. The “shrinkage” method used an adaptive shrinkage method (Schäfer & Strimmer, 2005b). The “glasso_0.01”, “glasso_0.001”, and “glasso_1e-4” methods used graphical LASSO with penalty parameters 10^-2^, 10^-3^, and 10^-4^ (Friedman, Hastie, & Tibshirani, 2008). Stronger regularization may lead to estimates that fit the data worse and also to worse-fitting knockoffs. Because setting the strength of shrinkage parameters is not fully understood in the context of knockoff construction, we tested a range of options empirically.

We evaluated the resulting knockoffs using three types of diagnostic. The first diagnostic determined how well each knockoff construction method preserved the data distribution. We concatenated the TF expression matrix with all TF expression knockoffs and jointly reduced to two dimensions via t-Stochastic Neighbor Embedding (t-SNE) (van der Maaten & Hinton, 2008) (Fig. S1A). Most methods appeared similar to the original data, but in the “permuted” method, the distribution of the knockoffs has very little overlap with the distribution of the original data. Based on this diagnostic, permuted knockoffs will not control FDR.

The second diagnostic is a technique based on k-nearest neighbors (KNN) that is sensitive to any violation of the key mathematical relationship to the data distribution that knockoffs must satisfy (Romano, Sesia, & Candès, 2019). For data with D variables and N observations, this test creates a matrix of size 2N by 2D, including data, knockoffs, and data randomly swapped with knockoffs. For any row of this matrix, the expected proportion of nearest neighbors that is swapped is 50%, and Romano et al. describe how to test this 50% proportion as a null hypothesis. Low p-values indicate evidence that knockoffs are invalid. Most knockoff generation methods failed this test, but the “sample”, “glasso_0.001”, and “glasso_1e-04” methods showed no evidence of poor fit (Fig. 2A). Based on this test, the “sample”, “glasso_0.001”, and “glasso_1e-04” model-X knockoff constructions would be expected to control FDR in testing for conditional independence.

**Figure 2.**
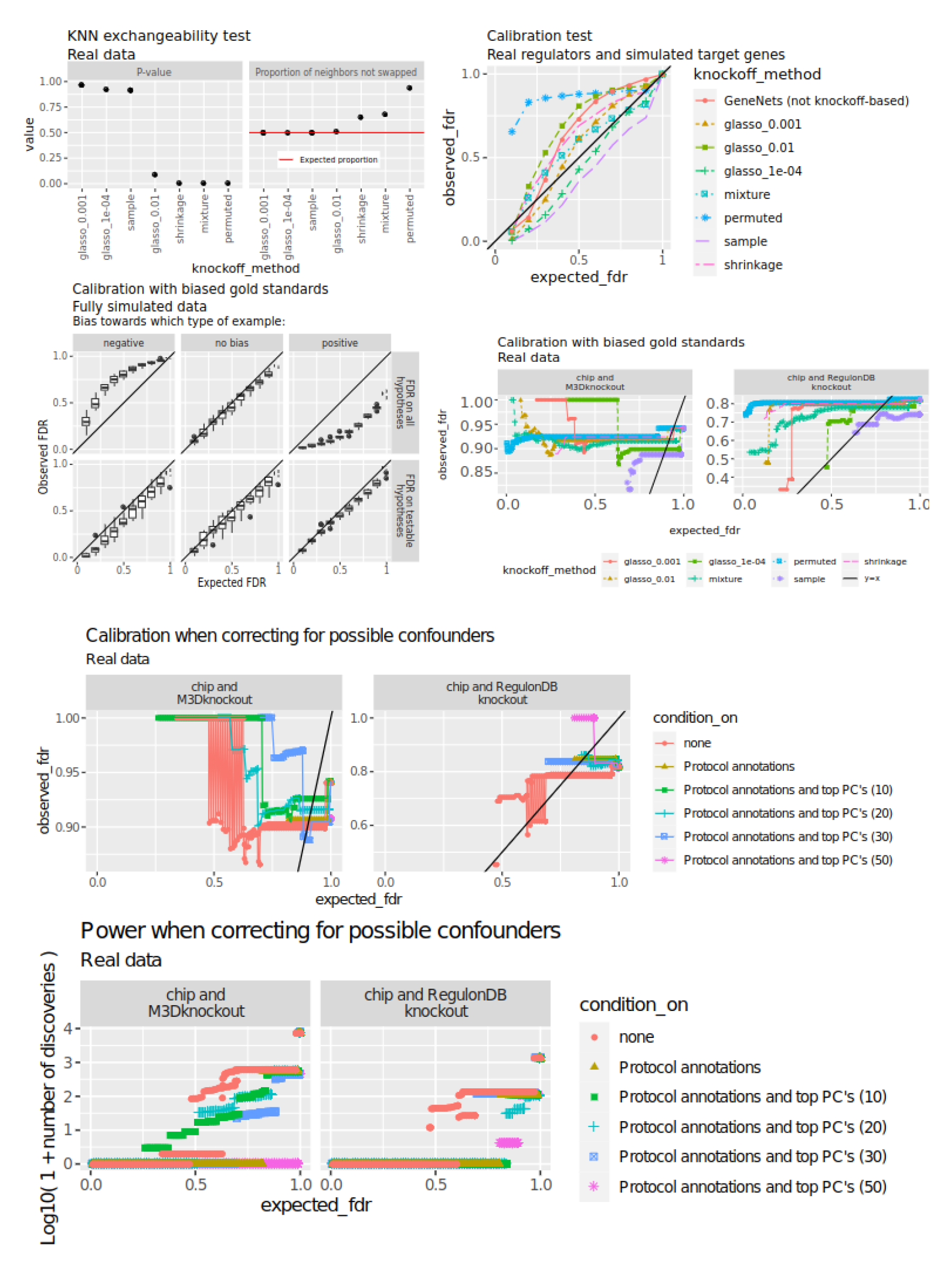
**Formal tests of conditional independence control FDR in testing conditional independence, but not in TRN inference.** This figure contains a mixture of real and simulated data. **A)** KNN-based exchangeability diagnostic from Romano et al. 2020 with k=20 and swapping all variables (“full”, not “partial” per Romano et al.). Low p-values (vertical axis) and high proportions of non-swapped neighbors (horizontal axis) indicate a poor fit. **B)** Expected and observed FDR of various knockoff generation methods and the GeneNet R package in selecting regulators for each of 1000 simulated *E. coli* target genes. Real regulator expression is used to simulate target genes. Each target genes is a step function applied to a single regulator. **C)** Expected FDR (from the knockoff filter) versus observed FDR (from the gold standards) on fully synthetic data using biased gold standards that contain excess positive or negative examples compared to the true proportion of positive or negative examples. The top row applies the final step of the knockoff filter to all hypotheses, while the bottom row applies the final step to testable hypotheses only. **D)** Comparison of expected FDR (from the knockoff filter) and observed FDR (from gold standards) across different gold standards and knockoff-generation methods, using real data. Ranges show 1.96 times the typical binomial standard error sqrt (p*(1-p) / n). **E)** Expected and observed FDR when correcting for labeled and unlabeled indicators of confounding. Knockoffs are constructed using the “glasso_1E-04” method conditioning on labeled perturbations, or on both labeled perturbations and the top (10, 20, 30, 50) principal components of the full expression matrix. Ranges show 1.96 times the typical binomial standard error sqrt (p*(1-p) / n). **F)** Power (number of discoveries) in the same analysis shown in panel E.

In the third diagnostic, we followed a commonly used simulation scheme that uses real TF expression and simulated target gene expression (Algorithm 1). Using real regulator data and simulated targets adequately tests the assumptions of model-X knockoff construction, which only requires modeling of regulators, and not targets. Using the same simulated targets, we also benchmarked the GeneNet R package, which controls FDR assuming the target gene is a linear function of its regulators (Schäfer & Strimmer, 2005a). GeneNet and most knockoff-based methods fail to control FDR, with permuted knockoffs performing worst. The “sample” and “glasso_1e-04” knockoff constructions control FDR in testing conditional independence (Figure 2B).

These three diagnostics characterized several attempts at FDR control in tests of conditional independence using real TF expression data. Based on the combined results, we conclude that the “sample” and “glasso_1e-04” knockoff constructions controlled FDR in testing conditional independence.

### Algorithm 1: Measuring FDR control with synthetic target genes

Input:

- TF expression *X* (matrix, N observations by G genes)
- Adesired FDR level *α*
- The number of simulations *J*
- A set of indices *i*_1_, *i*_2_, … *i_J_* indicating the true regulator in each simulation (integers between 1 and G)
- A set dose-response curves *f*_1_, *f*_2_, … *f_J_* dictating the true response to the regulator in each simulation

Procedure:

- For *j* = 1….*J*:

- Generate 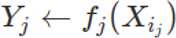. We set *f_j_*(*x*) = 1 if *x* > *mean*(*x*) and *f_j_* = 0 otherwise.
- Run the knockoff filter to obtain pairs (*i*, *j*) at FDR *α*.
- For *j* = 1….*J*:

- If *i_j_* is in the input, count (*i*,*j*) as correct.
- If *i_j_* is not in the input, count (*i*, *j*) as a false discovery.

### A modification to the knockoff filter allows empirical checks of FDR on incomplete gold standard data

In addition to tests on simulated target genes, tests on real target genes are desirable, because unlike simulated target genes, real data may not conform to the causal sufficiency assumption. A key obstacle to empirical tests of FDR control is incomplete or inaccurate gold standard data. Below, we will describe gold standards based on direct binding data (ChIP-chip, ChIP-seq, and ChIP-exo) and genetic perturbations followed by transcriptomics. Where the two sources disagree, edges will be annotated as unknown. The resulting gold standards should contain minimal errors, since each edge has two types of supporting evidence. However, these gold standards may contain disproportionately more positive or negative examples, because they include only the most confident conclusions and they focus on only a few genes. Because the base rate of positive examples does not match the network as a whole, naive checks of observed versus expected FDR are misleading.

To remove bias in FDR checks using incomplete gold standards, we partitioned the TF-target relationships from each gold standard into three sets: a set of positives P, a set of negatives N, and a set of unknowns U. Each hypothesis was considered testable if it was in P or N. We carried out the final step of the knockoff filter (the Selective SeqStep procedure (Barber & Candès, 2015)) on testable hypotheses or on all hypotheses using simulation studies. Focusing the analysis on testable hypotheses aligned the expected FDR from the knockoff filter with the observed FDR from the gold standard, whereas including all hypotheses failed to align the FDR (Fig 2C). This simulation suggests that the knockoff filter will control FDR against gold standards consisting of high-confidence positive and negative TF-target relationships, as long as the final step is applied to testable hypotheses only.

We developed two gold standards based on convergent results of distinct experimental designs. For a TF-target relationship to be included as a testable hypothesis we required concordant evidence from both ChIP data and genetic perturbation followed by transcriptomic analysis, rather than a replicated result between similar experiments (eg. multiple ChIP experiments). For one gold standard, we collected ChIP targets and knockout data from RegulonDB v10.9 (Santos-Zavaleta et al., 2019), and for the other we combined RegulonDB ChIP data with all genetic perturbation outcomes from the Many Microbe Microarrays Database (except where genetic effects were confounded by differences in growth medium). To check the reliability of these sources, we compared each dataset against the others and against RegulonDB v10.9 (Santos-Zavaleta et al., 2019), which is a manually curated collection supported by evidence from binding motif occurrences, binding assays, site mutation, or gene expression assays. We also compared against a small number of validation experiments from the Dream5 competition (Marbach et al., 2012). The various sources were well-supported by one another, except for RegulomeDB knockout data, which frequently did not support hypotheses from other sources and thus may be under-powered (Fig. S1B). These two gold standards contained 754 positive and 8496 negative examples across 6 TF’s, with each example having two types of concordant evidence. Combined with our method for testing FDR control on unbalanced gold standards, this resource provides a tractable method to check FDR control on a real TRN task.

### Conditional dependence does not imply direct regulation in the DREAM5 *E. coli* **expression data**

In the case of a TRN, causal sufficiency requires that all factors regulating transcription be observed; if causal sufficiency is met, then the observed and expected FDR should align when tested with real target genes as it did with simulated targets. We performed knockoff-based TRN inference, checked results on the testable hypotheses from the two gold standards. Greater excess FDR on calibration checks using simulated targets in the previous analysis (Fig. 2B) predicts greater excess FDR relative to gold standard data (Fig. 2D), and relative to gold standards, conditional independence testing via the knockoff filter outperforms permutation-based testing. The “sample” and “glasso_1e-04” methods, which successfully controlled FDR on simulated data (Fig 2B), also had the best FDR control of the included models (Fig 2D). However, the “glasso_1e-04” method failed to control FDR overall when applied to real target genes. The “sample” method displayed very low power on real data, with almost no q-values of testable hypotheses below 0.5, so we were unable to assess observed FDR for sets of hypotheses with low expected FDR.

As an example of how different FDR control methods can affect biological interpretation, consider the melibiose regulator MelR, which has in total 3 or 4 targets (Grainger et al., 2004; Wade, Belyaeva, Hyde, & Busby, 2000). Analyses using permuted knockoffs yielded 131 predicted targets of MelR. These discoveries were nominally at 1% FDR, but only one discovery (MelA) was a known target. The spurious targets detected by permutation-based FDR control span diverse biological functions, and if taken literally, these findings would massively revise the functional role of MelR. By contrast, the “glasso_1e-04” and “sample” methods do not discover any MelR targets at 1% FDR. This shows that using modern conditional independence tests can reduce FDR on a real TRN task, with meaningful improvement in interpretation.

Since FDR control with simulated targets does not translate to FDR control with real targets, this microarray dataset must not meet the causal sufficiency criterion. There are many possible reasons that the causal sufficiency assumption could fail. One possible reason is confounding by technical factors (Cote et al., 2022; Parsana et al., 2019). Another is exogenous perturbations: for example, repressor proteins can be activated by binding to a ligand, and this does not require altered mRNA levels (Semsey et al., 2013). We sought to address these possibilities with a combination of labeled perturbations present in the data and estimation of unobserved confounders via unsupervised machine learning.

To address possible confounding, we tested against gold standards while conditioning on labeled perturbations and principal components. Combined with the “glasso_1e-04” knockoff construction method, this approach effectively removed associations with all factors explicitly conditioned upon, producing very high q-values that indicate no evidence for conditional dependence (fig. S1C). Conditioning on labeled perturbations and principal components had little effect on the calibration relative to either gold standard or on the total number of discoveries (fig. 2E,F). However, power was low and very few discoveries contributed to the final benchmarks (fig. 2F). It remains unclear whether accounting for confounders in transcriptome data alone could mitigate the false discoveries driven by the causal sufficiency assumption, and furthermore, use of PCA as a proxy for unmeasured confounders may limit power by removing useful signal.

In principle, excess FDR on real target genes compared to simulated targets can have other causes aside from failure of the causal sufficiency assumption. Many TRN inference methods do not estimate the causal graph structure; instead, they infer a closely related undirected graph called the Markov random field (MRF) structure (Friedman et al., 2008), (Haury, Mordelet, Vera-Licona, & Vert, 2012) (Schäfer & Strimmer, 2005a), (Meinshausen & Bühlmann, 2010) (Kim, 2015), (Kotiang & Eslami, 2020), (Margolin et al., 2006; McDavid, Gottardo, Simon, & Drton, 2019)). Though the nodes of the MRF are identical to the nodes of the causal graph, the MRF has extra edges (Spirtes et al., 1993). Specifically, the neighbors of a node Y in the MRF consist of the parents, the children, and the spouses (parents of children) of node Y in the causal graph. This has been recognized as a complication by some causal structure learning methods (Pellet & Elisseeff, 2008). We accounted for this by treating all TF-TF edges as unknown and excluding them from the real-data calibration estimates. Thus, spousal relationships cannot explain the excess FDR we observe, and failure of the causal sufficiency assumption remains the likely culprit.

### Conditional dependence does not imply direct regulation in mouse skin RNA-seq data with paired chromatin state

Statistical assumptions that work or fail for TRN inference in *E. coli* may not work or fail the same way in eukaryotes (Marbach et al., 2012). Furthermore, modern multi-omic methods merge mRNA measurements with much more molecular information, and this may suffice to capture influences missed in mRNA data. In particular, genome-wide averages of activity near specific motifs may contain information about TF activity that is not present in transcript counts (Pemberton-Ross, Pachkov, & van Nimwegen, 2015). To evaluate knockoff filter FDR control on multi-omic data, we turned to a mouse skin and hair follicle dataset consisting of paired RNA and chromatin measurements on 34,774 single cells from female mice (S. Ma et al., 2020). We first applied our models to the RNA portion of the SHARE-seq data to demonstrate excess FDR, and then we incorporated the ATAC portion to check if excess FDR was reduced.

To reduce the effect of measurement error, we averaged the data across cells within 100 k-means clusters and discarded any cluster with <10 cells, producing 57 clusters. This is a reasonable method for separating biological and technical variation, since a similar approach has been shown to yield groups of cells that are consistent with an identical expression profile perturbed by multinomial measurement error (Baran et al., 2019). We generated permuted knockoffs and Gaussian knockoffs for the resulting TF expression matrix (57 clusters by 1,972 TF’s). Since there are more TF’s than expression profiles, Gaussian knockoffs could not be constructed using the “sample” method as done in the *E. coli* analyses; instead, a positive-definite optimal shrinkage estimator was used (Schäfer & Strimmer, 2005b). We did not attempt to fit Gaussian mixture models to the data, since there are only 57 observations and each cluster would require estimation of a mean parameter of dimension 1,972 as well as a 1,972 by 1,972 covariance matrix.

As in the *E. coli* analysis, we used two diagnostics to evaluate conditional independence tests prior to addressing questions of causality: simulated target genes and the swap-based KNN test. For simulated target genes, Gaussian knockoffs controlled FDR and permuted knockoffs did not (Fig. 3A). The KNN exchangeability diagnostic found no evidence against Gaussian knockoffs, but strong evidence for failure of permuted knockoffs, suggesting that genes in natural data are not all statistically independent (Fig. 3B). This demonstrates that permutation-based methods are unlikely to control FDR in TRN inference or in the simpler sub-task of conditional independence testing, but the knockoff filter can control FDR at least in conditional independence tests.

**Figure 3.**
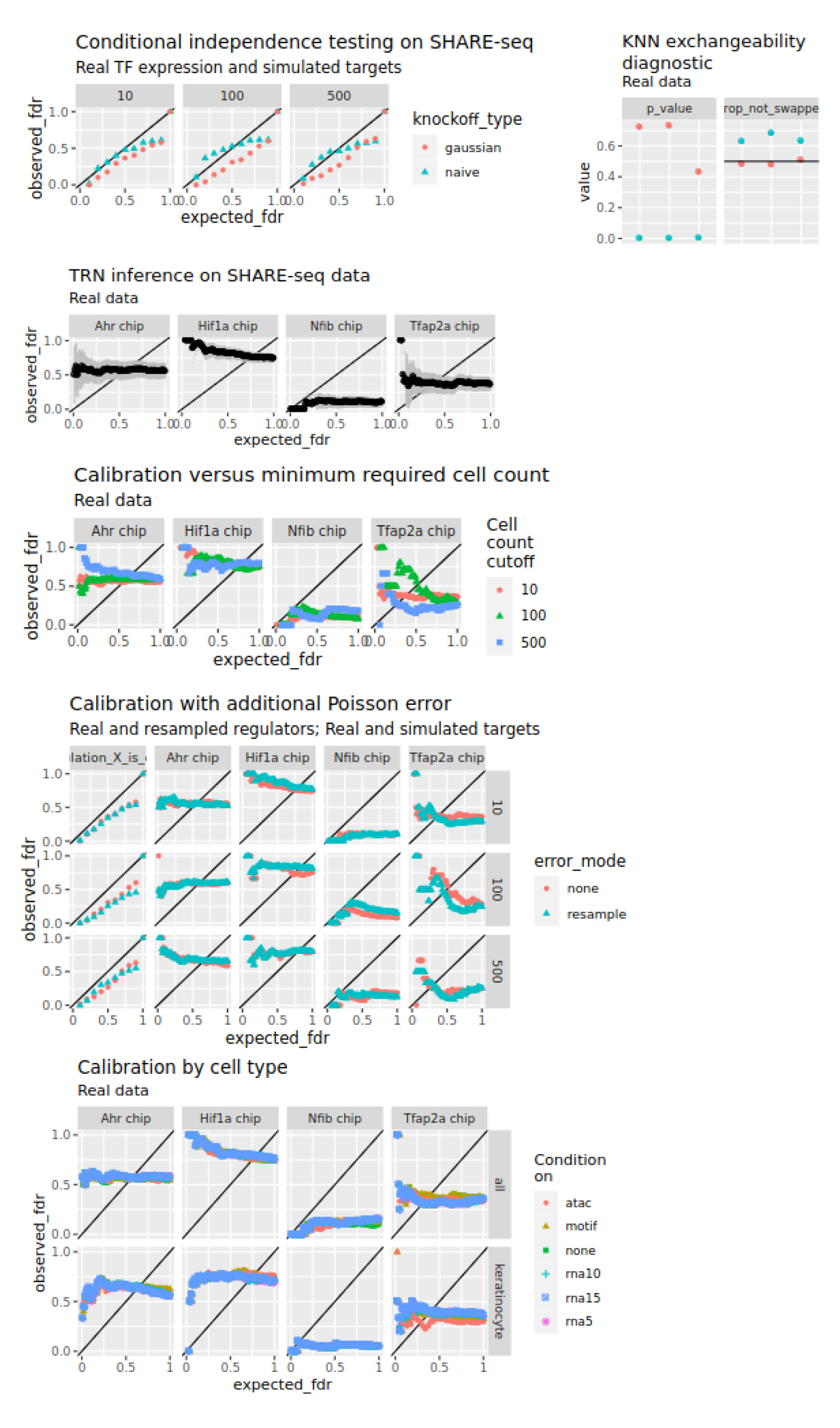
**Formal tests of conditional independence do not control false discovery rates with respect to ChIP-based gold standard data in a mouse skin network inference task, whether or not information on chromatin state is included.** This figure contains a mixture of real and simulated data. **A)** Error control in detecting regulators based on simulated target gene expression. Simulation uses real SHARE-seq TF expression data. Targets are simulated by applying a step function to a randomly chosen regulator. Knockoffs are constructed via independent permutation of each gene’s values (“permuted”) or Gaussian model-X knockoffs using an optimal shrinkage estimator (Schäfer & Strimmer, 2005b) for the covariance matrix. **B)** KNN exchangeability diagnostic results from Romano et al. with k=5 and swapping all variables (“full”, not “partial” per Romano et al.) (Romano et al., 2019). Low p-values (vertical axis) and high proportions of non-swapped neighbors (horizontal axis) indicate a poor fit. Results are shown for permuted and Gaussian knockoffs deployed on SHARE-seq TF expression data. **C)** Expected FDR from the knockoff filter versus observed FDR relative to ChIP-seq data when using the knockoff filter to infer regulators of each gene in the SHARE-seq data. Gaussian knockoffs are used. Error bars are 1.96 times the typical Binomial standard error. **D)** Variant of the experiment in panel C where clusters having fewer than the indicated number of cells are omitted. **E)** Variant of panels A (left column, simulated targets) and D (remaining columns) where covariates are contaminated with additional Poisson error prior to construction of knockoffs. **F)** Variant of panel C where knockoffs are constructed conditional on principal components of the SHARE-seq RNA matrix, principal components of the SHARE-seq ATAC matrix, or global activities of JASPAR mouse motifs in the ATAC matrix. The entire dataset (top row) or only keratinocyte-related cell types (bottom row) are used.

To test FDR on real gold standards, we selected four TF’s from ChIP-atlas with epidermal ChIP-seq data: NFIB, AHR, HIF1A, and TFAP2A (Oki et al., 2018). TFAP2A and HIF1A ChIP data were available in melanocyte cell lines and AHR and NFIB ChIP data were available in keratinocyte-related cell types. We used Gaussian knockoffs to infer regulators of all genes passing a minimum expression cutoff. ChIP-based results indicated poor enrichment and many false positives with highly confident results, with 37,824 findings at an FDR of 0.1 (Fig. 3C). This shows that conditional independence testing via knockoffs is not sufficient to control FDR in TRN inference on the RNA portion of the SHARE-seq data.

One potential explanation for this issue is measurement error: in a simple causal structure where A regulates B and B regulates C, then C and A are independent only given observations of B that are noiseless. To reduce the effect of measurement error, we increased the cutoff to 100 cells or 500 cells per cluster (38 and 28 clusters remained). Fewer discoveries were made (9,349 and 6,453 with q<0.1) but effects on enrichment were inconsistent (Fig. 3D). Increasing read depth in this way does not eliminate all measurement error, so we devised an independent method to estimate the degree to which measurement error increases false discoveries. We simulated measurement error starting from the cluster-aggregated data. Specifically, we resampled each TF expression count X_ij_ by replacing it with a Poisson draw having expectation equal to X_ij_. We constructed knockoffs based on the resampled TF expression. We tested the results on real target genes and on target genes that were simulated prior to resampling. Resampling caused slight deleterious effects in simulations, especially at high expected FDR, but had a weak effect on ChIP-seq benchmarks (Fig. 3E). Measurement error was not able to explain the degree of miscalibration we observed, implying a different type of missing information is driving the causal insufficiency that we observe.

Aside from transcript quantification errors, another driver of differences between conditional dependence and direct regulation is the inability of transcriptomics to directly measure TF activity. TF activity may be better captured by global motif accessibility, rather than the mRNA level of the TF (Balwierz et al., 2014; Madsen et al., 2018; C. Z. Ma & Brent, 2021). Therefore, we repeated our experiments while also conditioning on two types of global summaries of motif activity: the total fragments overlapping all occurrences of JASPAR 2018 mouse motifs (Khan et al., 2018) in the called peaks, and the top 10 principal components of the ATAC-seq count matrix. We also attempted to remove unmeasured confounding by conditioning on 5, 10, or 15 principal components of the gene expression matrix during knockoff construction. Results show hardly any difference (Fig. 3F), so in this case, even summaries of global chromatin accessibility do not contain enough additional information to approach causal sufficiency.

## Discussion

TRN inference methods are notorious for false positives (Diaz & Stumpf, 2022). For example, analyses using permuted genes as negative controls, which is sometimes claimed to control FDR in TRN inference (Chasman et al., 2019); (Kimura et al., 2020; Morgan et al., 2019), yielded 131 MelR targets spanning diverse biological functions. This conflicts with ChIP and perturbation experiments showing three or four targets of MelR, almost all located in the melibiose operon (Grainger et al., 2004; Wade et al., 2000). If MelR were not well-studied, follow-up experiments based on these findings could have wasted considerable resources. Such overconfidence is hard to detect and avoid in systems that lack ChIP or perturbation data, leaving users of TRN methods with no practical alternative but to distrust the results. Calibrated FDR control is an urgent unmet need for end users of TRN inference software.

Conditional independence testing and evaluation of causal sufficiency are two key steps towards successful FDR control in TRN inference. To advance this agenda, we departed from existing methods in two key ways. First, we controlled FDR in conditional hypothesis tests using model-X knockoffs. This does not require the strong linearity or independence assumptions of prior methods (Kim, 2015); (Chasman et al., 2019; Schäfer & Strimmer, 2005b); (Kimura et al., 2020; Morgan et al., 2019); (Mukhopadhyay & Chatterjee, 2007; Schäfer & Strimmer, 2005b). Though we recommend the knockoff filter be separately validated on each dataset it is applied to, we find that Gaussian knockoffs with regularized covariance estimates are a sensible initial choice for transcriptome data with low sample size and high dimension. Second, we used gold standards to evaluate FDR and calibration rather than area under the curve or other metrics. To detect a mismatch between simulated and real target genes, or between conditional dependence and direct regulation, known FDR is crucial, even if another method could obtain better AUPR against gold standard data without knowing the FDR. We demonstrated a way to evaluate FDR using incomplete gold standards, and we compiled incomplete *E. coli* gold standards with each example having concordant evidence from both perturbation transcriptomics and ChIP data.

In analyses of model-X knockoffs on mouse and *E. coli* datasets, FDR remained inflated. This was not explained by failure of conditional independence tests, which implies violation of causal sufficiency as the culprit. This is concordant with a recent report based on realistic simulations in which lack of causal sufficiency is expected to limit statistical TRN inference (Erbe et al. 2023). In light of these findings, existing results based on conditional independence alone should not be taken to reflect direct causal effects or direct binding, unless the underlying causal sufficiency assumption has been tested on the relevant datasets. Failure of causal sufficiency casts doubt on recent TRN work making explicit causal interpretations of conditional independence structure (Buschur, Chikina, & Benos, 2020; van Duin, Krautz, Rennie, & Andersson, 2022; Mohan, London, Fazel, Witten, & Lee, 2014; Qiu et al., 2020; Wang, Solus, Yang, & Uhler, 2017).

Where TF binding motifs are known, we expect motif analysis of open chromatin to substantially reduce false discoveries of TF-target interactions. Methods like Cicero (Pliner et al., 2018) often find motifs in enhancers, then pair enhancers with target genes via co-accessibility. Using co-accessibility to select upstream enhancers is similar to using co-expression to select upstream TF’s, and the knockoff filter could potentially be used for FDR control in enhancer-gene pairing, with the gold standard being chromatin conformation capture or expression quantitative trait locus data instead of ChIP-seq or TF knockouts. Pairing of enhancers with target genes is a crucial goal in the study of transcriptional regulation, with important consequences for interpreting genetic variants; this is a promising area of future study.

Other methods or technologies may yield measurements that reflect TF activity better than mRNA levels alone (Chung et al., 2021); (Chen et al., 2021; Qiu et al., 2020; Specht et al., 2021). Even given transcriptome data alone, methods like ARMADA (Pemberton-Ross et al., 2015) use genome-wide average motif activity, not mRNA levels, as a proxy for TF activity. These alternatives may come closer to satisfying the causal sufficiency assumption, but this will need to be empirically tested for each new system and data type. In the SHARE-seq skin example, we found the causal sufficiency assumption was violated even for TF mRNA plus global frequency in open chromatin of all JASPAR mouse motifs.

If measurements are never able to attain causal sufficiency, different analytical approaches may be needed. Causal structure inference methods not requiring causal sufficiency exist (Pellet & Elisseeff, 2008; Verny et al., 2017), and they have been deployed for TRN inference (Pellet & Elisseeff, 2008; Verny et al., 2017), but they lack FDR control. Equipping these methods with realistic and empirically validated guarantees on FDR control is an important area for future work.

## Conclusions

Lack of FDR control is a crucial bottleneck in biological interpretation of inferred TRN’s. Permutation tests do not control FDR in TRN inference or in the easier sub-task of conditional independence testing. Model-X knockoffs control FDR in conditional independence testing, but not in TRN inference. This mismatch occurs because the data used for TRN inference does not satisfy causal sufficiency. Methods controlling FDR in TRN inference must either explicitly check the assumption of causal sufficiency, or avoid it.

## Methods

### Code and data availability

All code used in this study will be made available at https://github.com/ekernf01/knockoffs_paper. All non-simulated data used in this study were publicly available when acquired, and will be made available as a collection via Zenodo at DOI 10.5281/zenodo.6573413.

### Knockoff filter usage

Knockoff construction is done via the R package `rlookc`, which is released along with this paper. The knockoff filter is applied using the same measure of variable importance throughout unless otherwise noted. It is the signed max lasso coefficient at entry (`stat.lasso_lambdasmax` from the R package `knockoff`) with one computational speedup: LASSO paths are fitted by glmnet with dfmax=21, corresponding to the assumption that no gene has over 20 direct regulators. In situations where FDR control is desired for a collection of discoveries from separate runs of the knockoff filter, for example across multiple *E. coli* target genes, FDR is estimated *after* pooling knockoff statistics.

### Speed and memory tests (Fig. S2AB)

Runtime was measured using the microbenchmark R package and peak memory usage was measured by the peakRAM R package.

### Threshold selection tests (Fig. S2C)

Covariates were simulated with the same mean, covariance, and sample size as the *E. coli* TF expression data. Knockoffs are constructed using the exact mean and covariance (not an estimate from the simulated dataset). Responses are set equal to the covariates, so each column has a single relevant feature. The knockoff filter is applied using the difference in linear model coefficients as the variable importance measure. Thresholds are selected separately for each target (“separate”) or using a single shared threshold (“merged”). FDR is calculated as the number of false discoveries across all targets divided by the number of discoveries over all targets.

### BEELINE benchmarking (Fig. 1)

Gaussian knockoffs were constructed using the sample mean and covariance. Gaussian mixture model parameters were inferred using mclust (Scrucca et al. 2016). We used 100 clusters, all having equal, spherical covariance. BoolODE does not separate production from decay, so RNA decay rates were inferred from RNA rates of change using piecewise quantile regression on RNA level. Self edges cannot be reliably inferred by our method and are ruled out a priori.

### E. coli datasets and gold standard processing

*E. coli* microarray data were downloaded from the DREAM5 challenge website at https://www.synapse.org/#!Synapse:syn2787211. The DREAM5 competition contains decoy genes with values chosen at random from the rest of the dataset (Cokelaer et al., 2015). These are absent from all gold standards, but we left them unchanged to facilitate comparison with previous work. They are easily distinguished from real genes by their low correlation with their knockoffs (Fig. S1D). *E. coli* transcriptional units were downloaded from the Biocyc smart table "All transcription units of E. coli K-12 substr. MG1655", available at https://biocyc.org/group?id=:ALL-TRANSCRIPTION-UNITS&orgid=ECOLI. Gold standard data were collected as follows.

- "dream5 validation”: interactions were manually extracted from Supplementary Data 7 of (Marbach et al., 2012).
- “M3Dknockout” includes all single-knockout samples and their controls from the DREAM5 training data, downloaded from https://www.synapse.org/#!Synapse:syn2787211. Experiments with aliased effects were not included; e.g. if the knockout was accompanied by a change in growth conditions relative to the controls. Any sample used in this gold standard was removed from the training data prior to knockoff construction whenever this gold standard was used for evaluation.
- "regulondb10_9" consists of manually curated regulatory interactions downloaded from https://regulondb.ccg.unam.mx/menu/download/full_version/files/10.9/regulonDB10.9_Data_Dist.tar.gz on 2022 Jan 28.
- "chip" and "regulonDB knockout": ChIP-based and knockout-based TF-target pairs were downloaded from RegulonDB version 10.9; a complete list of accessions is given in table S6. In *E. coli* ChIP data, IHF targets were regarded as targets of both IHF genes (ihfA and ihfB). MelR targets MelA and MelB were added manually, since they were missing despite having high-quality ChIP evidence (Grainger et al. 2005). ChIP-chip and ChIP-seq studies lacking loss-of-function controls were excluded to reduce risk of false positives (Waldminghaus & Skarstad, 2010); otherwise, all datasets listed were included.

*E. coli* targets are often determined at the level of a transcription unit, which may contain multiple genes (e.g. Nonaka et al. 2006, Kim et al. 2018). We thus augment *E. coli* ChIP and knockout-based gold standards to include any gene sharing a transcriptional unit with a target gene listed in the RegulonDB high-throughput downloads. For figures mentioning "chip and M3Dknockout" or "chip and RegulonDB_knockout", regulatory relationships consistent with both ChIP data and knockout data are treated as positive. Relationships missing from both are treated as negative. Additionally, the target and the regulator must each appear at least once in both datasets, or else the example is treated as unknown.

### E. coli analysis

*Knockoff filter.* For the 334 TF’s in the *E. coli* microarray data, knockoff features were constructed under multivariate Gaussian or Gaussian mixture model assumptions. When n>p, the semidefinite program implementation in the R package `knockoff` was used to determine optimal valid correlations of knockoff features with the original features. When p>n, a new method was used as described in Supplemental File S4. For mixture models, hard cluster assignments were set using the k-means clusters described below, and per-cluster covariance was estimated using the method for p>n.

Users simulating target variables to test reliability of Gaussian model-X knockoffs should be aware of a peculiar “double robustness” property: if Y is linear in X, FDR control will be maintained, even if X is wildly non-Gaussian (Huang & Janson, 2020). Therefore, Y should *not* be linear in X, or the diagnostic will never detect problems. After observing this phenomenon, we selected a family of unit step functions as the default for simulating Y in our software. Simulated target genes were constructed by selecting a single TF at random and simulating a target as I(X>m), where X contains the expression levels of the selected TF, m is the mean of X, and I() is an indicator function equal to 0 or 1. 1000 simulations were performed, and experiments cycled through 10 independently generated sets of knockoffs.

For each gene in turn, TF regulators were selected via the knockoff filter. To find regulators of TF’s, new knockoffs were created omitting the TF in question, and regulators were otherwise inferred in the same way. The efficient implementation described in Appendix 1 was used.

To adjust for confounders, knockoffs were computed after appending columns (features) to the TF expression matrix containing either non-genetic perturbations or non-genetic perturbations and the top principal components (Fig. 2E). The principal components were computed using the full expression matrix as input, scaled and centered. (These knockoffs thus violate the dictum to construct knockoffs without influence of the target variable, but the effect is to make the results more conservative.) Association with the confounding variables (Fig. S1C) was tested using the Pearson correlation as the measure of variable importance inside the knockoff filter.

T-SNE embeddings were computed using the R package `tsne` with default settings, using as input the 334 by 805 TF expression matrix concatenated with many 334 by 805 matrices of knockoffs (yielding a a 334 by 805*(k+1) matrix). K-means clusters were computed using the `kmeans` function from the R package `stats` with the entire expression matrix (TFs and non-TF genes) as input. Where multiple knockoff analyses are shown in the main figure, the t-SNEs in the supplement correspond to the analysis with no adjustment for confounders and no special handling of genetic perturbations.

Since the knockoff filter tests conditional independence, not the direction of causality, backwards edges confirmed by a given gold standard are marked as correct. To rule out spouses as a source of false positives (appearing in MRF structure but not gold standards), all TF-TF edges are marked as unknown, even if they appear to be ruled in or out by a given gold standard.

### Incomplete gold standard simulations (Fig. 2C)

Data are simulated, and knockoff statistics are computed, as in *Fig. S2C.* Gold standard positives (negatives) are marked unknown with 80% probability in the negative (positive) bias trials. Q-values are computed via Selective SeqStep (Barber & Candès, 2015) on all hypotheses (top row of figure), or only hypotheses that are testable with gold standard data (bottom row). Observed FDR is computed using gold standard data. Ten independent replicates are performed.

### Share-seq analysis

SHARE-seq skin count matrices were downloaded from GEO accessions GSM4156608 and GSM4156597 and reformatted as 10x-format HDF5 matrices using the DelayedArray and HDF5Array R packages. To successfully merge ATAC read counts with cell metadata, it was necessary to subtract 48 from the number in the final barcode associated with each cell in the count data. SHARE-seq data were briefly reanalyzed using the bioconductor packages scran, scater, and mbkmeans. Data were normalized by dividing by total counts per cell. 2000 highly variable genes were selected as input for PCA. 50 principal components were used as input for mbkmeans. Raw counts were summed within 100 clusters determined by mbkmeans. For keratinocyte-only experiments, existing cell-type annotations were used, and cells with the following labels were retained: ahighCD34+ bulge, alowCD34+ bulge, Basal, Hair Shaft-cuticle.cortex, Infundibulum, IRS, K6+ Bulge Companion Layer, Medulla, ORS, Spinous, TAC-1, TAC-2.

Pseudo-bulk expression profiles were normalized by dividing by total counts and multiplying by 1,000,000. Genes below 1CPM were excluded. Each gene was centered and scaled to have mean 0 and variance 1. Genes with constant expression were replaced with standard Gaussian random draws. Mouse TF’s and cofactors were downloaded from <u>AnimalTFDB 3.0</u> (Hu et al. 2019). Cofactors were used in addition to TFs since they can alter the effect of the TFs on downstream expression. Knockoffs were constructed for the centered, scaled TF expression matrix using the “permuted” method (permuting samples within each gene independently) or using the scalable Gaussian knockoff implementation in the function “createHighDimensionalKnockoffs’’ released in the `rlookc` package accompanying this paper. In cases where we test independence conditional on principal components of the ATAC or RNA data, these were computed using all genes/features, and they were concatenated onto the TF expression data prior to knockoff construction.

ChIP-seq data were downloaded from ChIP-atlas (Oki et al., 2018), counting all peaks for AHR, TFAP2A, NFIB, and HIF1A that fell within 10kb of a promoter as evidence that the respective TF regulates that gene. The following files were used:

http://dbarchive.biosciencedbc.jp/kyushu-u/mm10/target/Ahr.10.tsv
http://dbarchive.biosciencedbc.jp/kyushu-u/mm10/target/Hif1a.10.tsv
http://dbarchive.biosciencedbc.jp/kyushu-u/mm10/target/Nfib.10.tsv
http://dbarchive.biosciencedbc.jp/kyushu-u/mm10/target/Tfap2a.10.tsv

### Hardware and software used

All code used will be released prior to publication. Tests in figures 1 and S2 were run on a Dell XPS 13 with 8GB RAM and an Intel Core i5 processor. BoolODE was run in a virtual environment according to the maintainers’ instructions, with minimal changes made to export protein concentrations and RNA rates of change. BEELINE was run within a conda environment according to the maintainers’ instructions (https://github.com/Murali-group/Beeline). Minimal modifications were made in order to test multiple sets of parameters (https://github.com/Murali-group/Beeline/issues/59) and to benchmark directed and undirected FDR. *E. coli* and SHARE-seq experiments ran on Amazon Web Services EC2 t2.2xlarge instances based on the Ubuntu 18.04 image or on a Dell XPS15 running Ubuntu 20.04. Experiments used R version 4.1.2 and R package versions installed from either Bioconductor 3.14 or from MRAN’s archives from November 5, 2021. Seeds were set and package installation was automated for exact repeatability.

## Supporting information

Derivation of leave-one-out knockoffs

derivation of fast high-dimensional knockoffs

fdr control upon merging sets of discoveries

data downloaded

## Acknowledgements

Many thanks to Prashanthi Ravichandran, Da (Dan) Peng, Ashton Omdahl, and Josh Weinstock for helpful discussions, and again to Josh Weinstock for testing parts of the code. AB was funded by NIH grant R35GM139580. PC was funded by NIH grant R35GM124725.

## Author Contributions

E.K. conceived the study, derived knockoff filter speed-ups, performed data analysis, and created the figures. P.C. and A.B. supervised and funded the study. R.K. assisted in revising the manuscript. P.C. and A.B. guided choice of application datasets. E.K. wrote the manuscript with contributions from all authors.

---

**Figure S1.**
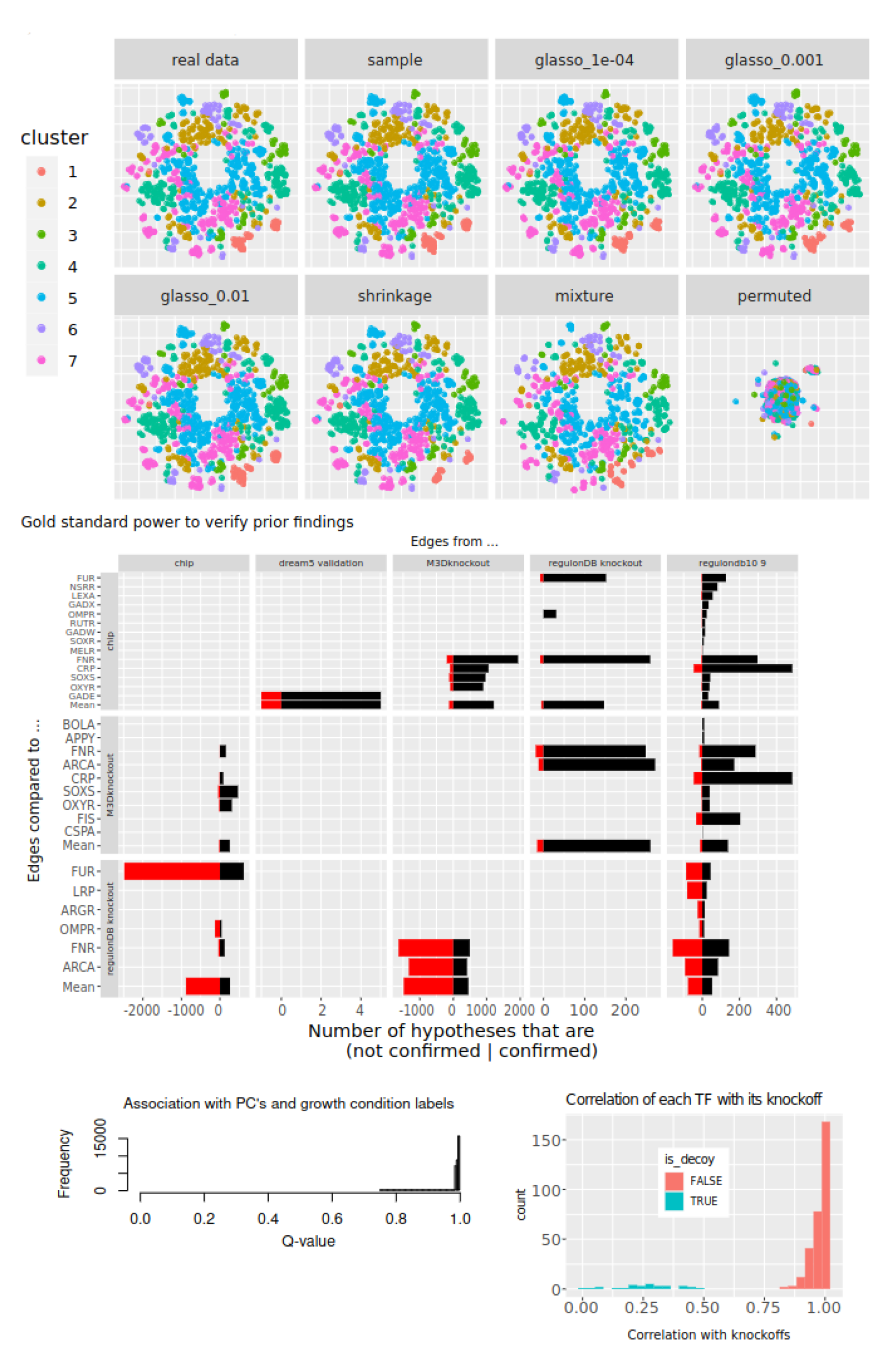
**Technical characteristics of knockoff construction on the *E. coli* TF expression dataset.** This figure contains only real data. **A)** T-SNE embeddings produced using TF expression and corresponding knockoff features. Samples are colored based on k-means cluster assignments, which were trained on the TF features. **B)** Comparison of gold standards. Hypotheses are extracted from the gold standard labeled in the top margin and checked to see if they are supported by the gold standard labeled in the left-hand margin. Hypotheses are omitted if they cannot be checked by the gold standard on the left, for instance if it is based on ChIP or knockout data and the regulator was never ChIPped or knocked out. **C)** Q-values for testing of conditional independence between each TF and each of the principal components or perturbation indicators that was explicitly conditioned on in Figure 2E. Knockoff generation method is “glasso_1E-04”. **D)** Correlation between variables and their knockoff copies. Gaussian knockoffs are generated for the 805 by 332 matrix of *E. coli* TF expression, using the sample covariance matrix.

**Figure S2.**
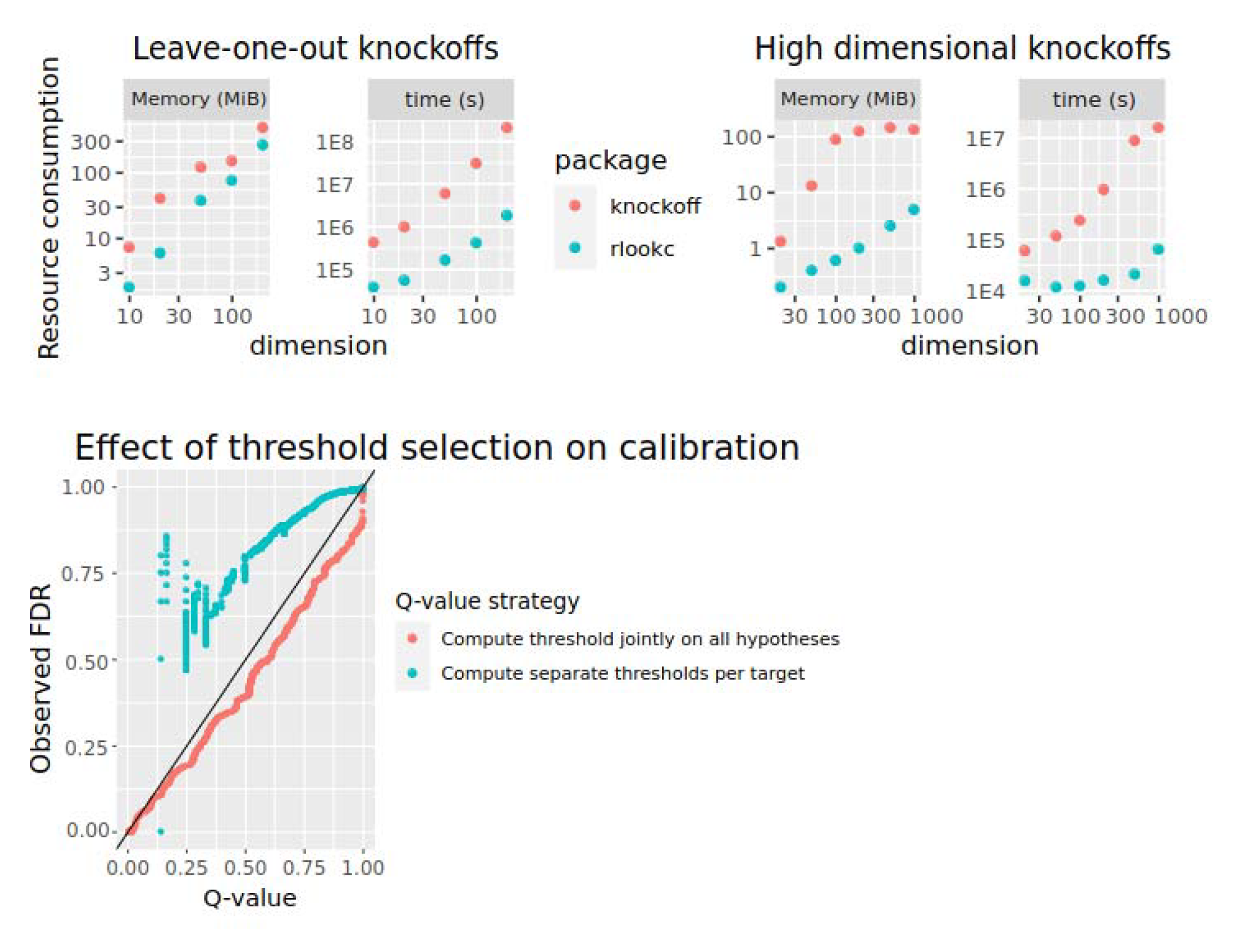
**Knockoff construction for transcriptome-scale data.** This figure contains only simulated data. **A)** Runtime and memory consumption for leave-one-out Gaussian knockoff construction with 1000 observations using our method and the reference implementation in the R package “knockoff”. **B)** Runtime and memory consumption for high-dimensional Gaussian knockoff construction with 10 observations using our method and the reference implementation in the R package “knockoff”. **C)** Threshold selection. Expected FDR from knockoff filter (x-axis) versus observed FDR (y-axis) in a simulated variable selection problem with 805 observations, 334 features, and 334 response variables. One trend line shows separate threshold selection for each target and the other shows a single threshold used across all targets.

## Supplementary info to be submitted

Supplementary info S1 (in this document): a user’s introduction knockoff filter concepts

Supplementary info S2 (in this document): performance enhancements included in our software

Supplementary info S3 (separate file): derivation of leave-one-out knockoffs

Supplementary info S4 (separate file): derivation of fast high-dimensional Gaussian knockoffs

Supplementary info S5 (separate file): rationale for choosing a single threshold across all target genes

Supplementary info S6 (separate file): table with high-throughput datasets downloaded from RegulonDB.

## Supplementary file S1

The knockoff filter (Barber & Candès, 2015; Candes, Fan, Janson, & Lv, 2018) is a framework for selecting a subset of features in a supervised machine learning model. The knockoff filter has been applied in genome-wide association studies and CRISPR-QTL studies; it has improved FDR control in cases where alternative methods fail, especially when the target variable has many inputs or when its functional relationship to the predictors is poorly understood (Sesia, Katsevich, Bates, Candès, & Sabatti, 2020) (Barry, Wang, Morris, Roeder, & Katsevich, 2021).

The principle underlying the knockoff filter is formal testing of *conditional independence*. Two variables Y and X are conditionally independent given a third set of variables S if P(Y|X,S) = P(Y|S). In other words, they are conditionally independent if X contains no new information about Y once S is known. The version of the knockoff filter we use accepts, as input: a desired FDR (Q); observations of a target (Y); observations of some features (X); and a distribution or probability model (P) that is assumed to have generated each observation in X. Due to the assumption that the model P generated X, this version of the knockoff filter is said to use *model-X knockoffs*. The knockoff filter returns a subset S(1), S(2), … S(k) of features in X where the FDR is below Q. The FDR is defined as the expected fraction of features in S where X_S(k)_ is independent of Y conditional on all of X except X_S(k)_. In other words, the knockoff filter returns a set of features that mostly contain non-redundant information about Y.

To use the knockoff filter, one must generate a carefully constructed negative control feature (a *knockoff*) for each feature in X. These knockoffs are not uniquely determined by P(X) but are heavily constrained by it. Different software implementations have enabled knockoff construction from different families of distributions (Sesia et al., 2020); (Romano et al., 2019), and the simplest method of constructing knockoffs is to predict each variable in turn from the other variables and knockoffs ((Barber & Candès, 2015; Candes et al., 2018)). Users can check via standard visualization methods whether a given set of knockoffs shares the topology and multimodal structure of their original data, and can also check validity of knockoffs via certain automated tests based on nearest neighbors (Romano et al., 2019). Recent work has shown that when an incorrect distribution is used, FDR control is not suddenly lost but rather declines gradually with an increase in the KL divergence to the true distribution (Barber et al., 2020; Zhou, Li, Zheng, & Li, 2022). However, if knockoffs appear inadequate, then the knockoff generation method should be revised.

Independently permuting the entries of each feature yields valid knockoffs, but only if the features are assumed to be independent under P(X) (Huang & Janson, 2020). This suggests that existing permutation-based methods for FDR-controlled TRN inference (Chasman et al., 2019); (Kimura et al., 2020; Morgan et al., 2019) will only work if all genes are statistically independent, which is an overly strong assumption. Throughout our experiments, we include permuted features as a baseline type of knockoff construction.

The conditional independence structure targeted by the knockoff filter is closely related to many TRN inference methods, including Scribe and ARACNE (Margolin et al., 2006; Qiu et al., 2020), and it is closely related to classic work on inference of causal mechanisms from observational data (Spirtes et al., 1993). The key insight of these methods is to disambiguate direct from indirect regulation. For example, in a simple pathway where A regulates transcription of B and B regulates transcription of C, then A and C will be independent conditional on B, correctly reflecting the absence of direct regulation. Certain additional assumptions are required for conditional independence to correctly infer causal structure. These are explained in classic texts on causal inference (Spirtes et al., 1993) and an accessible introduction is given by Scheines (Scheines, 1997). Pinpointing issues with these assumptions relative to known biology is a key focus of our work, and the knockoff filter is a crucial tool to generate findings about conditional independence structure at a known level of confidence.

## Supplementary file S2

To make our analysis methods more accessible, we have created an open-source R package called rlookc (for leave-<u>o</u>ne-<u>o</u>ut <u>k</u>nockoff <u>c</u>onstruction). The rlookc package speeds up computations, reduces memory requirements, and provides diagnostic tools for practical use. Compared to the existing R package *knockoff*, our software speeds up analyses that require leaving out one gene at a time and constructing knockoffs for the remaining genes. If done naively, these calculations scale with the fourth power of the number of genes; we reduce this to the cube, amounting to a 1000-fold speedup for a 1000-gene network (Supplementary file S3; Figure S2A). The software also generates fast knockoffs for datasets with many genes and few samples using much less time and RAM than the existing R package *knockoff* (Figure S2B and Supplementary file S4). It generates knockoffs for Gaussian mixture models, a simple and flexible class of distributions previously lacking an open-source software implementation (Gimenez et al., 2019; “Unable to find information for 13741696,” n.d.). We solve a technical issue related to combining sets of discoveries, which requires that a certain thresholding step be applied jointly to results on all target genes, rather than separately for each gene (Supplementary file S5; Fig S2C). Our software also includes tools for model assessment, such as a goodness-of-fit test that can detect individual outliers or inadequacy of knockoffs. These features lower barriers to entry for users hoping to apply the knockoff filter to functional genomics data. The software is available at https://github.com/ekernf01/rlookc.

