## Supplementary material for "Model-X knockoffs reveal data-dependent limits on regulatory network identification": Derivation of leave-one-out knockoffs

### Appendix 1: leave-one-out knockoff construction (LOOKC)

Network inference on N genes usually requires running N regression models, where each gene in turn is treated as the target. In this derivation, a method is developed for fast construction of Gaussian-X knockoffs when each variable is omitted in turn, so that we can condition on every variable except the one omitted.

To provide complete published documentation of our software, we include derivations of certain additional features not used in our study of TRN inference.

#### Ordinary knockoff construction

From Candes et al. (2016), Gaussian knockoffs are constructed such that the centered, scaled data $X$ and the knockoffs $\tilde{X}$ have joint covariance

$$G = \begin{bmatrix} \Sigma & \Sigma - S \\ \Sigma - S & \Sigma \end{bmatrix}$$

. This ensures the correct exchangeability properties that lead to proper FDR control. Since the mean is 0 and the distribution is Gaussian, this covariance matrix completely specifies the distribution. Here, $\Sigma$ is the covariance of $X$ or an estimate thereof. $S$ is a diagonal matrix that can be specified by the user. There is one constraint on $S$ (it must yield a positive-definite $G$), but otherwise $S$ can be chosen freely. The specific choice can affect the method's power, and existing software can determine a good option for $S$ that is compatible with the methods described herein.

Since $X$ is known but $\tilde{X}$ must be generated, a sample is drawn not from $Pr(\tilde{X}, X)$ but from $Pr(\tilde{X}|X)$. This distribution can be derived with standard techniques and is given in the model-X knockoffs paper. The exact formulas in terms of $X$, $S$, and $\Sigma$ are reproduced below as needed.

#### Reducing the cost

Generating knockoffs involves matrix operations of order $O(ND^2)$ and $O(D^3)$ where $D$ is the number of variables and $N$ the number of observations. In general, knockoffs must depend on $X$ but must not depend on $Y$, so whenever a new variable is treated as $Y$, the construction would need to be repeated with that variable left out. If done naively, this would add a factor of $D$ to the runtime (where $D$ is the number of variables).

Instead, it is possible to generate all leave-one-out knockoffs (LOOKs) within a constant factor of the original $O(ND^2 + D^3)$ computation time. The method is:

1. For a first approximation, generate knockoffs for $X$ and omit column $k$.

2. Update the knockoffs to remove the residual influence of column $k$ on the remaining variables.

The exact updates are derived below. They can be done by adding two rank-one matrices to the initial approximation.

**Derivation of updates**

- Without loss of generality, assume we wish to omit the final column of $X$ prior to knockoff generation, and call this variable $k$.
- Let $G$ denote the joint covariance of features and knockoffs as in Candes et al. (2016). Let $G_{-k}$ denote $G$ but omitting variables $k$ and $k+D$. Both rows and columns are omitted. $G_{-k}$ is never formed explicitly, but it is important mathematically because it specifies a joint covariance for the distribution of our leave-one-out knockoffs $Pr(\tilde{X}_{-k}, X_{-k})$. To obtain valid knockoffs, one requirement is that $G_{-k}$ must remain positive definite. This is satisfied because for a positive definite matrix, any principal submatrix is also positive definite. $G_{-k}$ also satisfies the knockoff exchangeability criterion. Thus, no change is needed in the choice of $S$.
- Let $M$ and $C$ be the mean and covariance of $Pr(\tilde{X}|X)$ with no variables omitted. From Candes et al. (2016),
  - $M = X - X\Sigma^{-1}S$
  - $C = 2S - S\Sigma^{-1}S$.
- Let $S_{-k}$, $\Sigma_{-k}$, $(\Sigma^{-1})_{-k}$, $X_{-k}$, $\tilde{X}_{-k}$, $M_{-k}$, and $C_{-k}$ denote the obvious matrices but with variable $k$ omitted. ($X_{-k}$ is the "first approximation" mentioned above.) For $S$, $\Sigma$, and $C$, omitting a variable means omitting the column and the row. For $X$ and $M$, only the column is omitted. For $(\Sigma^{-1})_{-k}$, the inverse is computed first, and row and column $k$ are omitted later. This implies

$$E[\tilde{X}_{-k}|X] = M_{-k} = X_{-k} - X_{-k}(\Sigma^{-1})_{-k}S_{-k}$$

  and

$$Cov[\tilde{X}_{-k}|X] = C_{-k} = 2S_{-k} - S_{-k}(\Sigma^{-1})_{-k}S_{-k}$$

  .

- Let $\tilde{M}$ and $\tilde{C}$ be the desired mean and covariance of the distribution we need to sample from: $Pr(\tilde{X}_{-k}|X_{-k})$. It yields almost the same result as $M_{-k}$ and $C_{-k}$, but note that variable $k$ must be omitted before computing the inverse of $\Sigma$, not after:
  - $\tilde{M} = E(\tilde{X}_{-k}|X_{-k}) = X_{-k} - X_{-k}(\Sigma_{-k})^{-1}S_{-k}$
  - $\tilde{C} = Cov(\tilde{X}_{-k}|X_{-k}) = 2S_{-k} - S_{-k}(\Sigma_{-k})^{-1}S_{-k}$.

A reasonable guess would be to pre-compute knockoffs with all variables and omit portions of them at each iteration. This method is not quite correct on its own, because

$$(\Sigma_{-k})^{-1} \neq (\Sigma^{-1})_{-k}$$

.

Another way to understand this is to note that

$$Pr(\tilde{X}_{-k}|X_{-k}) \neq Pr(\tilde{X}_{-k}|X)$$

. But, these initial guesses are very close, and they can be corrected efficiently, which we will now show by comparing $M_{-k}$ to $\tilde{M}$ and $C_{-k}$ to $\tilde{C}$. Before that comparison, there is one preliminary to discuss. Partition $\Sigma$ and $\Sigma^{-1}$ as

$$\Sigma = \begin{bmatrix} A & c^T \\ c & d \end{bmatrix}$$

and

$$\Sigma^{-1} = \begin{bmatrix} E & g^T \\ g & h \end{bmatrix}$$

. In general, $A^{-1} \neq E$, but this can be resolved with a standard rank-one update:

$$A^{-1} = E - g^T h^{-1} g$$

. This is useful in correcting both the mean and the covariance.

#### Mean

Without loss of generality, assume we are omitting the final variable, at index $k$. The mean can be partitioned to isolate the variable to be removed:

$$M = [M_{-k}|M_k] = [X_{-k}|x_k] - [X_{-k}|x_k] \begin{bmatrix} E & g^T \\ g & h \end{bmatrix} S$$

.

The relevant block is

$$M_{-k} = X_{-k} - (X_{-k}E + x_k g)S_{-k}$$

.

This is the mean of the naive procedure (generate knockoffs first, then omit). By contrast, we need the result as if variable $k$ were removed *before* knockoff generation:

$$\tilde{M} = X_{-k} - X_{-k}(\Sigma_{-k})^{-1}S_{-k} = X_{-k} - X_{-k}A^{-1}S_{-k}$$

. The necessary correction is of rank 1. It is:

$$\begin{aligned}
\tilde{M} - M_{-k} &= -X_{-k}A^{-1}S_{-k} + (X_{-k}E + x_k g)S_{-k} \\
&= -X_{-k}ES_{-k} + X_{-k}g^T h^{-1} g S_{-k} + X_{-k}ES_{-k} + x_k g S_{-k} \\
&= X_{-k}g^T h^{-1} g S_{-k} + x_k g S_{-k} \\
&= (X_{-k}g^T h^{-1} + x_k) g S_{-k}
\end{aligned}$$

.

#### Covariance

For the covariance, the desired matrix (again removing variable $k$ *before* generating knockoffs) is

$$\begin{aligned}
\tilde{C} &= 2S_{-k} - S_{-k}(\Sigma_{-k})^{-1}S_{-k} \\
&= 2S_{-k} - S_{-k}A^{-1}S_{-k} \\
&= 2S_{-k} - S_{-k}ES_{-k} + S_{-k}g^T h^{-1} g S_{-k} \\
&= C_{-k} + S_{-k}g^T h^{-1} g S_{-k}
\end{aligned}$$

.

Thus, the covariance of the precomputed knockoffs can be corrected by adding a random vector $S_{-k}g^T h^{-1/2} z_n$ where $z_n \sim N(0, 1)$. This must be done $N$ times, once per observation in $X$.

These derivations have been implemented in our R package `rlookc` and successfully tested for correctness against the reference implementation in the R package `knockoff`.

#### Testing groups of variables

In a dataset with a correlated set of variables $\ell$ in $X$, it may be impossible to distinguish among the different options, yet it may be clear that at least one of them is in the active set. In this scenario it is desirable to test the null hypothesis

$$Y \perp\!\!\!\perp X_\ell | X_{-\ell}$$

(where indexing by $-\ell$ denotes omission of the whole set). In Sesia et al. (2020), model-X knockoffs are extended to composite hypotheses of this type. Their framework assumes variables are partitioned into (disjoint) groups $\ell_1, ...\ell_L$. Error control is similar to the un-grouped knockoff framework, but the exchangeability criterion

$$swap(B, X\tilde{X}) =^D X\tilde{X}$$

no longer needs to be met for all sets of variables $B$. Rather, exchangeability is only required for swaps where grouped variables stay together, i.e. $B$ is the union of any of the $\ell$'s.

A method for constructing such knockoffs is described in Sesia et al. (2018), but it only applies to a specific HMM used on genotype data. Constructing grouped model-X knockoffs for Gaussian $X$ is discussed in Katsevich and Sabatti (2019) as an extension of prior work done under the more restrictive assumptions of fixed-X knockoffs (Dai and Barber 2016). The method is reasonably simple: the diagonal matrix $S$ can be replaced with a block-diagonal matrix where the blocks correspond to the variable groups. As before, the only other constraint on $S$ is that $G$ must remain positive definite. Dai and Barber (2016) give a fast, simple scheme for choosing $S$.

Given knockoffs obeying the correct exchangeability criterion, test statistics may be constructed arbitrarily as long as they are symmetric under the null. For example, the maximum (or mean) importance measure within each group can be subtracted from the maximum (or mean) over the corresponding knockoff importance measures, or the test could use the likelihood ratio

$$\log \frac{Pr(Y|X_{-\ell}, \tilde{X})}{Pr(Y|X, \tilde{X}_{-\ell})}$$

, or LASSO-based methods could use grouped LASSO.

Grouped hypothesis tests are implemented in `rlookc` and unit-tested successfully, with checks for error control, power, and positive definiteness of $G$.

#### Groups of variables and leave-one-out knockoffs

The computational cost of sampling group knockoffs is similar to the cost of sampling individual knockoffs. For efficient leave-one-out knockoffs, the strategy outlined above applies with slight modifications. Partition $S$ as

$$S = \begin{bmatrix} S_{-k} & S_{k,-k} \\ S_{-k,k} & S_k \end{bmatrix}$$

.

Since $S_{-k,k}$ is no longer 0,

$$M_{-k} = X_{-k} - (X_{-k}E + x_k g)S_{-k} - (X_{-k}g^T + x_k h)S_{k,-k}$$

. The terms on the left are as above, but the rightmost term is new. The desired quantity has the same formula as before, though $S_{-k}$ may not be diagonal:

$$\tilde{M} = X_{-k} - X_{-k}(\Sigma_{-k})^{-1}S_{-k} = X_{-k} - X_{-k}A^{-1}S_{-k}$$

. The necessary correction is still of rank 1, and the algebra strongly resembles the case above.

$$\begin{aligned}
\tilde{M} - M_{-k} &= -X_{-k}A^{-1}S_{-k} + (X_{-k}E + x_k g)S_{-k} + (X_{-k}g^T + x_k h)S_{k,-k} \\
&= -X_{-k}ES_{-k} + X_{-k}g^T h^{-1} gS_{-k} + X_{-k}ES_{-k} + x_k gS_{-k} + (X_{-k}g^T + x_k h)S_{k,-k} \\
&= X_{-k}g^T h^{-1} gS_{-k} + x_k gS_{-k} + (X_{-k}g^T + x_k h)S_{k,-k} \\
&= (X_{-k}g^T h^{-1} + x_k)gS_{-k} + (X_{-k}g^T + x_k h)S_{k,-k} \\
&= (X_{-k}g^T h^{-1} + x_k)gS_{-k} + (X_{-k}g^T h^{-1} + x_k)hS_{k,-k} \\
&= (X_{-k}g^T h^{-1} + x_k)(gS_{-k} + hS_{k,-k})
\end{aligned}$$

.

Similarly, the last three terms ($\equiv R$) are new in

$$\begin{aligned}
C_{-k} =&2S_{-k} - [S_{-k}|S_{k,-k}] \times \Sigma^{-1} \times [S_{-k}|S_{k,-k}]^T \\
=&2S_{-k} - (S_{-k}ES_{-k} + S_{-k,k}gS_{-k} + S_{-k}gS_{k,-k} + S_{k,-k}hS_{k,-k}) \\
\equiv&2S_{-k} - S_{-k}ES_{-k} - R
\end{aligned}$$

, and the correction becomes

$$\begin{aligned}
\tilde{C} &= 2S_{-k} - S_{-k}(\Sigma_{-k})^{-1}S_{-k} \\
&= 2S_{-k} - S_{-k}A^{-1}S_{-k} \\
&= 2S_{-k} - S_{-k}ES_{-k} - R + R + S_{-k}g^T h^{-1} gS_{-k} \\
&= C_{-k} + R + S_{-k}g^T h^{-1} gS_{-k} \\
&= C_{-k} + S_{-k,k}gS_{-k} + S_{-k}gS_{k,-k} + S_{k,-k}hS_{k,-k} + S_{-k}g^T h^{-1} gS_{-k} \\
&= C_{-k} + bf + f^T b^T + bhb^T + f^T h^{-1} f \\
&= C_{-k} + (h^{1/2}b + h^{-1/2}f^T) \times (h^{1/2}b^T + h^{-1/2}f)
\end{aligned}$$

. For brevity, we have used some new names above: $b \equiv S_{-k,k}$ and $f \equiv gS_{-k}$. To convert a Gaussian random vector with covariance $C_{-k}$ into one with covariance $\tilde{C}$, it suffices to add a standard Gaussian times the square root of the increment, which is

$$h^{1/2}b + h^{-1/2}f^T = h^{1/2}S_{-k,k} + h^{-1/2}S_{-k}g^T$$

.

These results reduce to the updates for non-grouped LOOKs whenever $S_{k,-k} = 0$. Grouped LOOKs are successfully unit tested against the slower reference implementation for mean, covariance, and correlation with held-out variables.

**Gaussian mixture knockoffs and the efficient leave-one-out knockoffs (LOOKs)**

Compared to Gaussian knockoffs, Gaussian mixture models can provide more flexibility and extend the applicability of this framework. If an observation $X_i$ is drawn from a mixture of $J$ Gaussians with PDF $\sum_{j=1}^{J} \pi_j N(\mu_j, \Sigma_j)$, then Gimenez et al. (2019) showed that a joint density for $x_i$ and a knockoff could be $\sum_{j=1}^{J} \pi_j N(\mu_j, G_j)$, where $G_j$ is defined separately for each cluster as above. If $z_i$ is the (latent) cluster chosen for $x_i$, we can generate a knockoff by choosing a cluster $z_i$ at random from $P(z_i|x_i)$ and drawing knockoffs from $P(\tilde{x}_i|x_i, z_i)$; Gimenez et al. (2019) show that this preserves the necessary exchangeability property.

For mixture-model leave-one-out knockoffs (LOOKs), it is necessary to sample $z_i$ from $P(z_i|x_{i,-k})$, meaning almost the same posterior but without knowledge of one coordinate. Fast methods for doing this are outlined in the next paragraph. Then $\tilde{x}_{i,-k}$ can be drawn from $P(\tilde{x}_{i,-k}|x_{i,-k}, z_i)$ as described above. Beware: cluster assignments may vary across leave-one-out iterations even within the same observation, and the low-rank updates should not be applied to a knockoff observation sampled from the wrong cluster. It is thus necessary to maintain multiple initial guesses for the knockoffs, one per cluster, and always select the correct one to update.

To sample $z_i$ from $P(z_i|x_{i,-k})$, suppose the estimates of the cluster proportions $P(Z = z)$ remain unchanged. From Bayes' Theorem,

$$P(Z = z|X_{-k}) = \frac{P(X_{-k}|Z = z)P(Z = z)}{\sum_z P(X_{-k}|Z = z)P(Z = z)}$$

. If cluster $z$ has mean $\mu$ and covariance $\Sigma$, then computation is dominated by the cost of

$$P(X_{-k}|Z = z) = \frac{1}{\sqrt{\det(2\pi\Sigma_{-k})}} \exp[\frac{-(x_{-k} - \mu_{-k})^T (\Sigma_{-k})^{-1} (x_{-k} - \mu_{-k})}{2}]$$

.

The term inside the exponent can be cheaply obtained from the rank-one correction used earlier:

$$(\Sigma_{-k})^{-1} = A^{-1} = E - g^T h^{-1} g$$

. The determinant can be computed from a Cholesky decomposition: if $\Sigma = LL^T$ and $L$ is triangular, then $\det(\Sigma) = \det(L)\det(L) = (\prod_k \ell_k)^2$. One way to update the determinant cheaply is via the Cholesky factors. If

$$\begin{bmatrix} \Sigma_{-k} & \Sigma_{k,-k} \\ \Sigma_{-k,k} & \Sigma_{k,k} \end{bmatrix} = \begin{bmatrix} L_{-k}^T & L_{-k,k}^T \\ 0 & L_{k,k} \end{bmatrix} \begin{bmatrix} L_{-k} & 0 \\ L_{-k,k} & L_{k,k} \end{bmatrix}$$

, then

$$\Sigma_{-k} = L_{-k}^T L_{-k} + L_{-k,k}^T L_{-k,k}$$

. By a well-known matrix determinant lemma,

$$\det(\Sigma_{-k}) = \det(L_{-k}^T L_{-k}) \det(1 + L_{-k,k}(L_{-k}^T L_{-k})^{-1} L_{-k,k}^T)$$

. Given a precomputed $L$ of size $D$, this update can be computed in $O(D^2)$ time via forward and backward substitution, compared to $O(D^3)$ if done naively.

Gaussian mixture model knockoffs are implemented and unit-tested in our software, but leave-one-out Gaussian mixture model knockoffs are not yet implemented at time of writing.

Gimenez, Jaime Roquero, Amirata Ghorbani, and James Zou. "Knockoffs for the mass: new feature importance statistics with false discovery guarantees." The 22nd International Conference on Artificial Intelligence and Statistics. PMLR, 2019.
