## Supplementary material for "Model-X knockoffs reveal data-dependent limits on regulatory network identification": derivation of fast high-dimensional knockoffs

### Appendix 3: high-dimensional Gaussian knockoffs

RNA-seq and ATAC-seq commonly measure 20,000 to hundreds of thousands of features. The original model-X knockoffs paper includes a GWAS demo with 71,145 SNPs. But, the SNPs are distributed over 23 chromosomes with each chromosome treated separately, and the biggest matrix operations are on the order of a 10k by 10k block. Since RNA-seq and ATAC-seq can far exceed the 10,000 variables included in the original demonstration of model-X knockoffs, we envision the need for more computationally efficient high-dimensional Gaussian knockoff generation. Here, we develop an efficient method for Gaussian knockoff generation in settings with p>>n, where the dominant cost is that of a singular value decomposition (SVD).

##### Estimating covariance

The sample covariance matrix will be singular and a poor estimate of the true covariance. We instead begin with the optimal shrinkage method of Schaefer and Strimmer, which yields a positive definite estimate as well as better mean squared error than the sample covariance.

We assume throughout that X has mean 0, variance 1 for each feature. If needed, knockoffs can be constructed on centered, scaled data and then transformed back to match the original mean and scale. Let $S$, $G$, and $\Sigma$ denote the same matrices that were used in the leave-one-out derivations in appendix 1.

##### Setting $S$

The first task in generating Gaussian knockoffs for large $p$: it is hard to find $S$ such that

$$\begin{bmatrix} \Sigma & \Sigma - S \\ \Sigma - S & \Sigma \end{bmatrix}$$

is positive definite. The memory requirements of the reference implementation appear to scale with the square of the dimension. But, the shrinkage estimator above suggests a much easier way to obtain a valid $S$. It returns an estimate of the form

$$\Sigma = (1 - \lambda)R + \lambda I$$

where $R$ is the sample covariance matrix and $\lambda$ is determined from the data by `corpcor::estimate.lambda`. Since a sufficient condition for $G$ to be positive definite is $2\Sigma - S$ to be positive definite, $S$ can be set to $2\rho\lambda I$ for $0 < \rho < 1$, and then

$$2\Sigma - S = 2((1-\lambda)R + \lambda I) - 2\rho\lambda I = 2(1-\lambda)R + 2(1-\rho)\lambda I$$

which is positive definite.

##### Sampling knockoffs

It is useful to discuss certain computational tricks prior to the rest of the derivation. The sample covariance $R$ is $\frac{1}{n-1}X^TX$. Let $UTV$ be a full SVD of $X$ with singular values $t_i$. Then the column $i$ of $V$ is an eigenvector of many related matrices:

- $R$ with eigenvalue $\frac{t_i^2}{n-1}$
- $\Sigma$ with eigenvalue $(1-\lambda)\frac{t_i^2}{n-1} + \lambda$
- $\Sigma^{-1}$ with eigenvalue $[(1-\lambda)\frac{t_i^2}{n-1} + \lambda]^{-1}$
- $[2S - S\Sigma^{-1}S]^{1/2}$ with eigenvalue $[\Big[4\rho\lambda - 4\rho^2\lambda^2[(1-\lambda)\frac{t_i^2}{n-1} + \lambda]^{-1}\Big]^{1/2}$

Thus, the product $MZ$ where $M$ is any of these matrices can be computed as:

$$MZ = VV^TMVV^TZ = Vf(T)V^TZ$$

where $f(T)$ returns a diagonal matrix with the appropriate transformation applied piecewise to the eigenvalues in $T$.

For $i > n$, $t_i = 0$, so anything orthogonal to the top $n$ columns of $V$ has the same eigenvalue. Partition $V$ accordingly as $[V_X|V_\perp]$. The product can be written

$$MZ = V_Xf(T)V_X^TZ + f(0)V_\perp V_\perp^TZ$$

. This can be computed with just $O(np)$ memory because

$$V_\perp V_\perp^TZ = Z - V_X(V_X^TZ)$$

.

##### Knockoff mean

The knockoffs must be created with mean

$$X - X\Sigma^{-1}S$$

. The reference implementation in the R package `knockoff` forms $\Sigma$ and computes the inverse explicitly, which incurs prohibitive $O(p^2)$ memory requirements. Instead, we apply the third bullet point above, with $f$ chosen for $\Sigma^{-1}$.

##### Knockoff covariance

The knockoffs must be created with covariance $C = 2S - S\Sigma^{-1}S$. To do this, the reference implementation in the R package `knockoff` performs an inverse and a Cholesky factorization, both of them having prohibitive $O(p^2)$ memory requirements. Instead, we use the fourth bullet point above to efficiently multiply by the square root of $2S - S\Sigma^{-1}S$.
