## Supplementary material for "Model-X knockoffs reveal data-dependent limits on regulatory network identification": fdr control upon merging sets of discoveries

### Appendix 2: FDR control of unions of sets

*This material is adapted from one author's comments on a public question and answer forum, licensed under a Creative Commons CC-by-SA 4.0 license and available here.*

The knockoff filter involves a choice of threshold, which is varied based on the user's desired FDR and which is applied to a set of intermediate statistics produced as part of the knockoff filter. Here we motivate the choice to use a single threshold across all target genes, rather than running the whole knockoff filter procedure for each gene separately and later combining the discoveries.

Consider merging two disjoint sets of discoveries, each generated by a method that controls FDR at level $\alpha$. Let $a$ and $b$ be the number of false discoveries in the first and second set respectively. Let $c$ and $d$ be the total number of discoveries. Let $\lambda = \frac{c}{c+d}$. "FDP" will refer to the false discovery proportion observed in the data (with the convention FDP=0 for cases with no discoveries), and "FDR" will mean $E[FDP]$.

The combined FDP is a convex combination of the individual FDP's.

$$\begin{aligned} \frac{a+b}{c+d} &= \frac{a}{c+d} + \frac{b}{c+d} \\ &= \frac{c}{c+d}\frac{a}{c} + \frac{d}{c+d}\frac{b}{d} \\ &= \lambda\frac{a}{c} + (1-\lambda)\frac{b}{d} \end{aligned}$$

It is tempting to conclude

$$E[\frac{a+b}{c+d}] = \lambda E[\frac{a}{c}] + (1-\lambda)E[\frac{b}{d}] = \alpha$$

where $\alpha$ is the FDR of the two input sets, but since $\lambda$ is random and not independent of $a, b, c, d$, this equality does not hold in general. A counterexample can be constructed as follows. Let $\alpha = 0.5$, let $c - a = 1$, let $b = 1$, let $d = 2$, and let $a$ equal 0 or 100 with 50% probability. Then, $E[\frac{a}{c}]$ is in fact slightly less than 50%, but $\frac{a+b}{c+d}$ is 1/3 or 101/103 with equal probability, and its expected value exceeds 0.5. When the number of true hypotheses is limited, as we expect in a sparse gene regulatory network, the mixture proportion $\lambda$ is dominated by sets that happen to yield more false discoveries. Thus, in practice, FDR is inflated when measured across the combined set. In our implementation, we do not control FDR independently for selecting regulators of each target gene. Rather, we choose a threshold jointly across all target genes. Joint choice of threshold is not mathematically guaranteed to our knowledge, but in simulations, it seems to resolve this issue.
